# Life stage and vaccination shape the gut microbiome of hatchery-reared Atlantic salmon (*Salmo salar*)

**DOI:** 10.1101/2023.02.21.529474

**Authors:** Kara J. Andres, Bingdi Liu, Lauren E. Johnson, Kevin L. Kapuscinski, Ashley H. Moerke, Fangqiong Ling, Jason H. Knouft

## Abstract

Microbiomes play an essential role in promoting host health and fitness, but the factors affecting variation in gut microbiomes among individuals are not fully understood. Investigating the microbiome under different conditions is needed to link gut microbiomes to host physiology and potentially design manipulations to improve rearing success of captive species. In this study, we characterized the gut microbiomes of Atlantic salmon (*Salmo salar*) in individuals at different life stages, vaccination status, and hatchery origin. Microbiomes differed between age-0 sub-adults and adults, with sub-adults exhibiting higher diversity and more similar communities when compared to adults. We also found that vaccines against bacterial kidney disease reduced gut microbial diversity within individual sub-adult salmon, resulting in dissimilar gut microbial communities among individuals. The diversity and structure of microbiomes did not differ between groups of adults that were reared in two different hatcheries and sampled from the wild. Sub-adults, particularly unvaccinated sub-adults, displayed a strong core microbiome present in the majority of individuals. Our results suggest that life stage and vaccination status are essential factors in the gut microbiome development of salmon. Conditions experienced during early life stages appear to have a strong influence on the microbiome, but differences among individuals at early life stages may be lost due to environmental factors experienced later in life. The plasticity of the microbiome throughout the life of individuals may have important implications for understanding host health, with potential applications for improving the rearing and reintroduction success of the ecologically and economically important Atlantic salmon.

**IMPORTANCE:** The Atlantic salmon (*Salmo salar*) is a globally important fisheries and aquaculture species, but the factors affecting gut microbiomes of hatchery-reared fish are not fully understood. Our study explores the influence of life stage, vaccination status, and hatchery origin on the composition and structure of the Atlantic salmon gut microbiome. We found that life stage is an important driver of gut microbiome diversity, likely driven by differences in habitat and diet.

Vaccination against bacterial kidney disease led to marked declines in gut microbial diversity within individuals, resulting in highly distinct gut microbial communities among individuals. Hatchery origin did not have a strong influence on adult Atlantic salmon captured from the wild. These findings suggest that life stage and vaccination drive variation in Atlantic salmon microbiomes, but the stability and long-term implications of such variation on host health should be considered in future microbiome research.

## INTRODUCTION

The assemblage of microbes found within the gastrointestinal tract of host species, collectively termed the gut microbiome, can affect host phenotype and fitness through traits associated with fecundity, body weight, behavior, nutritional uptake, and disease resistance (1– 5). Understanding the link between gut microbiome and host health is of interest to conservation biologists and fisheries managers who may be able to leverage microbiome manipulations— through diet, pre- and pro-biotics, antibiotics, and/or vaccines—to enhance rearing and reintroduction success of commercially important, threatened, and endangered species (5–9). For instance, administering probiotic microorganisms has been used as a means to optimize immunity, enhance growth rates, and protect against pathogens in salmonids, a family of fishes with high economic value in aquaculture and commercial and recreational fisheries around the globe (10–12). However, despite a growing understanding of the important role of the gut microbiome in host health, the complex, dynamic, and potentially interacting factors shaping the structure and composition of microbial communities are poorly understood, particularly in non-model hosts. Determining the factors shaping variation in host microbiomes is therefore a requisite step in understanding how microbiomes contribute to host physiology, as well as how this relationship may be manipulated to mitigate disease and promote host health.

Conservation biologists, aquaculturists, and fisheries managers are particularly interested in improving the rearing and stocking success of Atlantic salmon (*Salmo salar*), a species that is endangered within much of its native range, but also of global importance in the aquaculture industry and supports important fisheries within the Laurentian Great Lakes (hereafter Great Lakes). These generalist predators can also stabilize aquatic food webs in Great Lakes communities that have a long and complicated history of anthropogenic impacts including habitat loss, overexploitation, invasive species, and pollution (13). Native to Lake Ontario and the north Atlantic Ocean, Atlantic salmon were extirpated from the Great Lakes by the end of the 19^th^ century as a result of dam construction, deforestation, overfishing, and pollution (14, 15).

The first hatchery to begin captive rearing and reintroducing Atlantic salmon into the lakes was established in 1866 (15), and since then over 1 million Atlantic salmon have been introduced across the Great Lakes. Although natural reproduction has been observed in the St. Marys River, Michigan (16), there is no evidence that recruitment to adulthood is sufficient to sustain a population. Fisheries are therefore dependent on stocking hatchery-reared Atlantic salmon, but stocking programs have shown variable success and often poor return rates (0–2%) (15, 17–20). Given that the stocking success of hatchery-reared individuals may be linked to the gut microbiome (21, 22), understanding how different environmental and host-associated factors shape the microbiome may be essential for improving hatchery success.

The primary factors governing the gut microbiome are broadly associated with characteristics of the host, environment, and diet (23), and hatchery conditions such as water quality and source, food supply, and environmental bacterial communities may therefore have a strong influence on Atlantic salmon microbiomes. Hatcheries may rear fish in recirculating or flow-through water systems, or in cages in open water, and the composition of the microbiome tends to differ among these rearing conditions (24, 25). Vaccinations administered to reduce diseases in hatcheries may also affect the gut microbiome of fishes through induced immunological responses (26), although the interactions between vaccination, immune response, and gut microbiome in hatcheries remain relatively understudied (27, 28). The specific diet provided to hatchery-reared individuals may also have a large influence on the gut microbiome (29), and signatures of hatchery dietary conditions may persist long after individuals are released into the wild (22). However, while hatchery conditions during early life stages can have large implications for the assemblage of gut microbiota and host health (30, 31), the gut microbiome is also extremely dynamic and may rapidly respond to changes in environmental or dietary conditions throughout the course of an individual’s life (32).

In this study, we used 16S rRNA gene metabarcoding to explore how gut microbial communities in Atlantic salmon vary among individuals of different life stages, vaccination status, and hatchery origins. First, we investigated general microbial community differences between sub-adult (age-0) and adult salmon. Previous research has shown that sub-adult salmon have lower microbial richness and reduced among-individual variation in microbial assemblages when compared to adult individuals (33). Next, we compared the microbial communities of sub- adult salmon that were vaccinated against bacterial kidney disease (BKD) to those that were not vaccinated. BKD is a chronic disease caused by the bacterium *Renibacterium salmoninarum* from the Corynebacteriaceae family that affects salmonid species around the world. The disease has no implications for human health but can cause up to 40% mortality in Atlantic salmon (34, 35). The BKD vaccine is a live vaccine and can be delivered by intraperitoneal injection, intramuscular injection, or oral administration. Although shown to significantly increase survival of Atlantic salmon in experimental trials (36), no work has investigated how BKD vaccines impact the gut microbiota in fish populations. Finally, we investigated whether hatchery origin of adult salmon produced a lasting impact on gut microbiome diversity and structure after adults lived in the wild for an extended period of time. We explored group comparisons through an in-depth examination of gut microbiomes, including characterization of alpha- and beta diversity as well as identifying core communities and biomarkers associated with each group.

## METHODS

### Salmon rearing and collection

In this study, we compared the microbial communities across 20 sub-adult (age-0) Atlantic salmon (10 BKD-vaccinated and 10 unvaccinated) and 18 adult Atlantic salmon (8 and 10 individuals originating from two different hatcheries; Figure S1). Sub-adult salmon were collected from Lake Superior State University’s (LSSU) Center for Freshwater Research and Education (CFRE) Fish Hatchery when they were approximately 10 months of age with a mean total length of 161.5 mm and weight of 44.1 g (Table S1). Sub-adults reared in this facility resulted from artificial spawning of approximately 100 pairs of returning adult Atlantic salmon collected from the St. Marys River, Chippewa County, MI, USA in fall 2019. Adult salmon stocked in previous years were collected in November 2019 and held in artificial raceways for gamete collection. Fertilized eggs were treated with 0.75% saline, 2 ppm (mg/L) erythromycin, 100 ppm (mg/L) iodine, and 1,000 ppm (mg/L) thiamine. Age-0 Atlantic salmon were moved into fiberglass fry raceways in late February 2020 and were reared in filtered (bag and UV filters) and heated St. Marys River water until June 2020, at which time they were transferred into large raceways (fiberglass or concrete) and reared in ambient river water until harvest for this study (7 and 11 November 2020). Vaccination to prevent BKD or to reduce the effects of the disease was administered to sub-adult Atlantic salmon via their feed at the LSSU hatchery during 4–7 November 2020. Sampled individuals were euthanized by personnel at the LSSU hatchery with an overdose of buffered MS-222.

Adult Atlantic salmon collected in this study were initially stocked in the St. Marys River originating from either the LSSU hatchery or the Michigan Department of Natural Resources’ (DNR) Platte River State Fish Hatchery. Artificial spawning for both hatcheries took place at the LSSU hatchery (with propagation and rearing methods described above), and eyed embryos were transferred to the DNR hatchery. Embryos and age-0 fish at the LSSU hatchery were reared as described above, and fish were reared in flow-through, fiberglass raceway systems supplied with untreated water from the St. Marys River from age 5 months until the time of stocking (approximately 17 months). Fish at the DNR hatchery were reared in concrete raceways supplied with UV-treated, sand-filtered, temperature-controlled water from a spring-fed pond until an age of about 6 months and were then reared on ambient Brundage Creek water until the time of stocking. Fish from both hatcheries were stocked in the St. Marys River and recaptured approximately 16 months later after living in, presumably, the St. Marys River and Lake Huron.

Adults were collected from the St. Marys River in November 2020 using short-set gill nets (15.2 m x 3.4 m) with a 10.2 cm mesh. Nets were directly observed while fishing and pulled immediately upon capture of a salmon. Upon removal from the net, each fish was measured for total length and weight, examined to determine sex, identified to hatchery origin based on the presence of specific fin clips, and euthanized with an overdose of buffered MS-222. The gastrointestinal tract of each individual was removed at LSSU by clamping the anterior and posterior ends and using scissors to disconnect the GI tract. All tissue was immediately placed in a falcon tube or plastic bag and stored in a -80°C freezer.

### Sample processing and DNA extraction

Frozen age-0 individuals and the gastrointestinal tracts of adult individuals were shipped overnight on dry ice from LSSU and received at Saint Louis University on November 20, 2020. All samples were immediately stored at -80°C. We emptied the gut contents of thawed fish specimens and gastrointestinal tracts into sterile petri dishes using disposable scalpels and steam-sterilized forceps and homogenized the contents in a sterile phosphate-buffered saline (PBS) solution using a fixed-speed rotator. Homogenized contents were passed through a sterile 70-µm cell strainer (Fisherbrand™) to remove large debris, and the pass-through suspension was made into aliquots for DNA extraction.

We extracted DNA from the gut content PBS mixture for each sample using bead beating, enzymatic treatments, and a phenol-chloroform extraction method (37). Briefly, 450 µl of DNA extraction buffer and 200 µl of sample were added to a microcentrifuge tube with 0.1 g of 0.1 mm glass beads. After bead-beating and cooling on ice, we added an enzymatic cocktail and 10% SDS to each tube, which was then vortexed and incubated at 37°C. This was followed by another incubation at 60°C with 10% CTAB and 5 M NaCl. We added 600 µl of phenol-chloroform-isoamyl alcohol (pH ∼8) and centrifuged each sample. The supernatant was washed with 600 µl chloroform-isoamyl alcohol twice and incubated at -20°C with sodium acetate, linear acrylamide and isopropanol. The DNA precipitate was then washed three times with 70% ethanol after centrifuging and then air-dried. Last, the DNA pellet was dissolved with biological grade molecular water at 55°C and stored at -20°C. DNA samples were purified with the Wizard® DNA Clean-up System and DNA concentrations were quantified using AccuBlue® High Sensitivity dsDNA Quantitation Kit. Four reagent negative controls were included alongside all gut content samples.

### 16S rRNA gene amplicon sequencing

We prepared sequencing libraries using a two-step PCR protocol (38). First, the 16S rRNA genes were amplified by polymerase chain reaction (PCR) with universal primers 515F (5’-GTGCCAGCMGCCGCGGTAA-3’) and 806R (5’-GGACTACHVGGGTWTCTAAT-3’) targeting the V4 hypervariable region of most bacterial and archaeal 16S rRNA genes (39). We visually inspected all PCR products by electrophoresis on a 1% agarose gel to confirm amplicon size. To minimize potential amplification biases, each sample was amplified in four separate reactions and PCR products were combined (38). We purified the products of first-step PCR using the Wizard® SV Gel and PCR Clean-Up System and used the purified products as templates for a second-step PCR in which unique barcode pairs were attached to each sample.

Each sample was amplified with two pairs of barcodes to generate two technical replicates in Illumina sequencing. We quantified PCR products using the AccuBlue® High Sensitivity dsDNA Quantitation Kit and submitted the samples for Illumina Miseq paired-end sequencing (2 × 250 bp) at the Center for Genome Sciences & Systems Biology at Washington University in St. Louis. Samples with fewer than 10,000 reads were re-sequenced and combined with reads from the first sequencing run.

### Bioinformatic processing

Demultiplexed DNA sequences were imported to the QIIME2 platform to perform the bioinformatic analyses (40). To achieve a quality score above 30, both forward and reverse reads were trimmed at 20 bp and forward reads were truncated at 118 bases. We then used the DADA2 algorithm to denoise sequencing data, remote putative chimeric sequences, and generate amplicon sequence variants (ASVs) (41). We performed taxonomic classification on the ASVs within QIIME2 using a multinomial naive-Bayes classifier trained against the SILVA-138 database (42, 43). The outputs of this pipeline, including the total number of sequence reads for each ASV in each sample and the taxonomic classification for all detected ASVs, were imported into R using the qiime2R package and converted to a phyloseq object (44, 45).

ASVs with fewer than three reads across all samples and ASVs originating from an unknown domain or Eukarya domain were removed (0.254% and 0.001% of the reads across all samples, respectively). Three out of four negative control replicates had more than 50 reads, and reads in these negative controls, as well as negative controls from other recent experiments conducted in the laboratory, were used to identify potential contaminant ASVs using the decontam package with the ‘prevalence’ method and threshold set to 0.5 (46). A total of eight ASVs were identified as contaminants with 14.39% relative abundance across all samples and were subsequently removed. At this stage, two samples were removed for failing to meet a minimum quality threshold of 10,000 reads.

To examine technical robustness, we calculated the Pearson correlation between technical replicates of the same sample based on relative read abundance, and all technical replicates resulted in *r* > 0.98 (Table S1), indicating high technical reproducibility. We therefore merged the reads of technical replicates for each sample. All samples were rarified by subsampling to 10,254 reads based on the minimum number of reads across all non-redundant samples.

### Data analysis

Analyses were conducted in R 4.1.2 (47) and results were visualized using ggplot2 (48). To identify differences in alpha diversity among compared groups (i.e., life stage: adults vs. sub-adults, sub-adult vaccination status: BKD-vaccinated vs. unvaccinated, and adult hatchery origin: LSSU vs. DNR), we calculated richness (total observed ASVs, Shannon diversity, and Pielou’s evenness) for each sample using package vegan (49, 50). Differences in alpha diversity among groups were evaluated by a non-parametric Kruskal-Wallis H test. All p-values were Bonferroni-corrected to account for multiple comparisons. We quantified the Kruskal-Wallis effect size difference between groups using package rstatix (51), where η^2^ < 0.06 indicates a small effect, η^2^ = 0.06–0.14 indicates a moderate effect, and η^2^ > 0.14 indicates a large effect.

To evaluate microbiome diversity among samples (beta diversity), we calculated Bray-Curtis dissimilarities between samples based on the ASV relative abundances [BC_*ij*_ = (∑ | x_*ij*_ - x_*ik*_|)/∑ (x_*ij*_ + x_*ik*_), where x_*ij*_ and x_*ik*_ are relative abundances of ASV *i* in samples *j* and *k*]. We performed a Principal Coordinates Analysis (PCoA) on the Bray-Curtis dissimilarity matrix using the ‘ordinate’ function in phyloseq and the ‘pcoa’ function in ape (49, 50). Differences in Bray-Curtis dissimilarities among groups were evaluated by permutational multivariate analysis of variance (PERMANOVA) using the function ‘adonis2’ and analysis of multivariate homogeneity of group dispersions (PERMDISP) using ‘betadisper’ from vegan.

To better understand the specific microbiota driving differences in microbiome composition among groups, we identified core microbiota and used linear discriminant analysis (LDA) to identify microbial biomarkers that distinguish communities from one another. We define core microbiota as those with high prevalence, or those that present in ≥ 80% of the samples in a particular group. The prevalence and the mean relative abundance of microbiota were calculated at the ASV, genus, and family levels for each group being compared. The LDA effect size (LEfSe) was performed using package lefser (52). Briefly, LEfSe examines if features from two sample groups are significantly different in abundances by first screening for taxa that reject a null hypothesis of equal average abundance, and then screening by effect sizes. As a first screen, differentially abundant features were screened for by passing a Kruskal-Wallis test with alpha = 0.05. Next, LDA was performed to screen for differentially abundant taxa by primary LDA scores. The primary LDA score of a taxon was calculated by averaging the differences between actual group means with the differences between group means along the first linear discriminant axis. The output of the analysis is a list of features with log-10 transformed LDA scores larger than 2, and these microbiome biomarkers were identified at family and genus level for the compared groups.

## RESULTS

Thirty-six salmon gut microbiota samples were analyzed, resulting in a total of 1,973,000 reads and 1,181 ASVs. Following rarefaction to 10,254 sequences per sample, a total of 369,144 reads and 1,105 ASVs remained.

Sub-adult gut microbiomes were dominated by *Firmicutes* (87.4% relative abundance), followed by *Proteobacteria* (10.0%) and *Actinobacteria* (1.9%; Figure 1; Table S2). In contrast, adult gut microbiomes were dominated by *Proteobacteria* (76.5%), with the next most common phyla being *Firmicutes* (18.7%), then *Actinobacteria* (1.4%). Vaccinated sub-adults exhibited a much larger proportion of *Proteobacteria* (20.5%) than unvaccinated sub-adults (0.5%).

**Figure 1.**
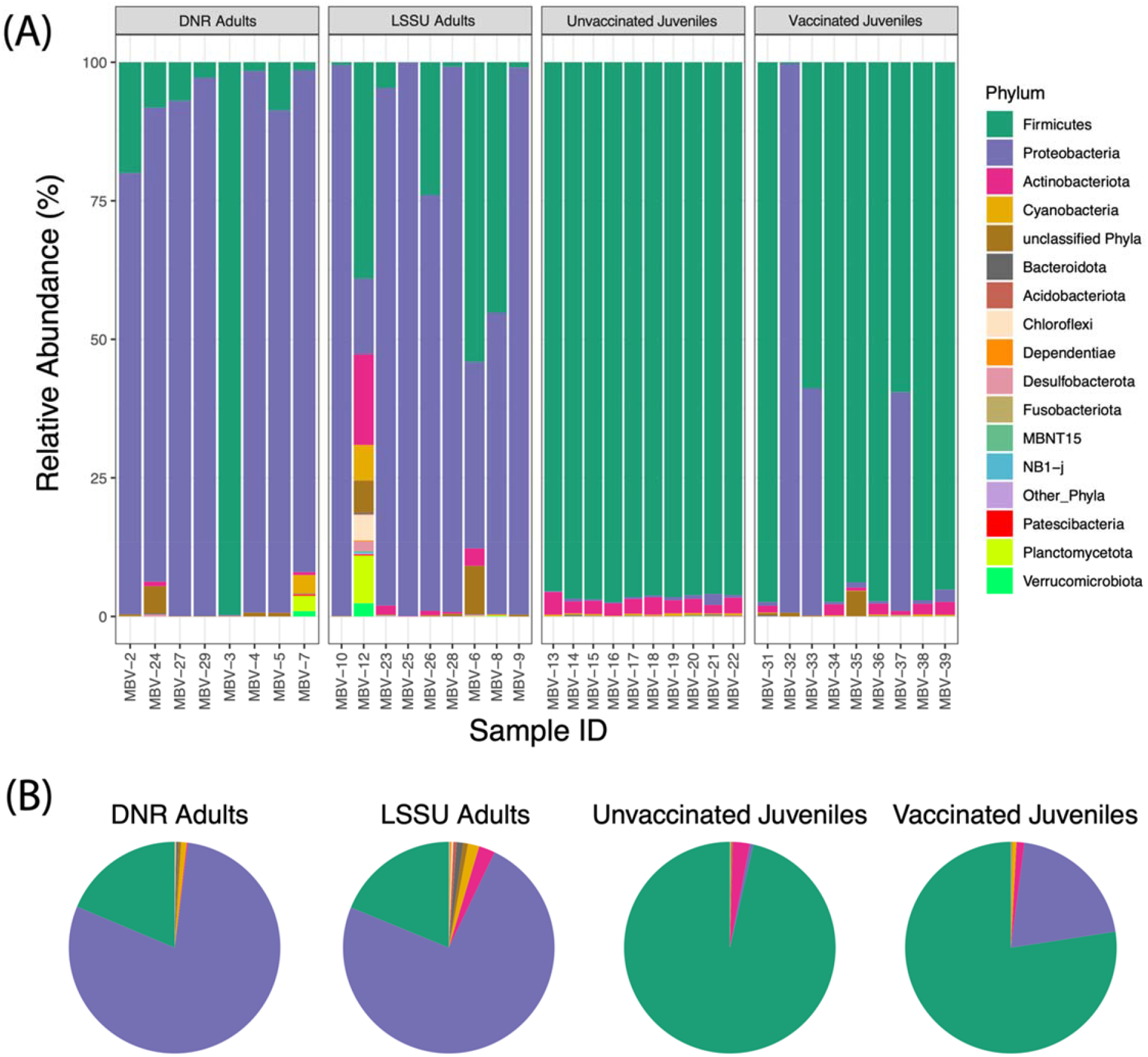
Phylum-level taxonomic compositions of Atlantic salmon gut microbiota. (A) Relative abundance based on read counts for each individual sample, grouped by host factors: adult hatchery origin and sub-adult BKD vaccination status. Phyla with relative abundance below 0.1% were binned as “other phyla”. (B) Average relative abundance of abundant phyla in adults from two different hatcheries and sub-adults that were either BKD-vaccinated or unvaccinated.

### Microbial alpha diversity

Sub-adults exhibited higher alpha-diversity than adults across all diversity indices (Kruskal-Wallis p_adjusted_ ≤ 0.001 for observed ASV richness, Shannon index, and Pielou’s evenness; Figure 2A; Table 1). BKD-vaccinated sub-adults exhibited lower ASV richness (p_adjusted_ = 0.007) and Shannon diversity (p_adjusted_ = 0.027) than unvaccinated sub-adults (Figure 2B; Table 1). However, Pielou’s evenness did not differ among vaccinated and unvaccinated sub-adults. We did not detect any differences in gut microbiome alpha-diversity metrics in adults originating from the different hatcheries (i.e., LSSU vs. DNR; Figure 2C; Table 1).

**Table 1.** Kruskal–Wallis comparisons of alpha diversity statistics for gut microbiomes of individuals differing in life stage [sub-adults (*n* = 19) vs. adults (*n* = 17)], sub-adult BKD vaccination [unvaccinated (*n* = 10) vs. vaccinated (*n* = 9)], and adult hatchery origin [LSSU (*n* = 9) vs. DNR (*n* = 8)]. P-values were Bonferroni-corrected for multiple comparisons and bold values indicate statistical differences at alpha = 0.05.

| | α-diversity | Sub-adults | Adults | $\chi^2$ | p-value |
| --- | --- | --- | --- | --- | --- |
| Life stage | ASV richness | 160.9 | 58.8 | 12.60 | **0.001** |
|  | Shannon index | 3.33 | 1.11 | 16.06 | **< 0.001** |
|  | Pielou's evenness | 0.66 | 0.28 | 16.84 | **< 0.001** |
| | α-diversity | Unvaccinated | Vaccinated | $\chi^2$ | p-value |
| Sub-adult vaccination status | ASV richness | 200.0 | 117.4 | 9.38 | **0.007** |
|  | Shannon index | 3.73 | 2.88 | 6.83 | **0.027** |
|  | Pielou's evenness | 0.71 | 0.61 | 1.71 | 0.574 |
| | α-diversity | LSSU | DNR | $\chi^2$ | p-value |
| Adult hatchery origin | ASV richness | 69.3 | 46.9 | 0.08 | 0.773 |
|  | Shannon index | 1.40 | 0.78 | 0.75 | 0.387 |
|  | Pielou's evenness | 0.33 | 0.22 | 0.93 | 0.336 |

**Figure 2.**
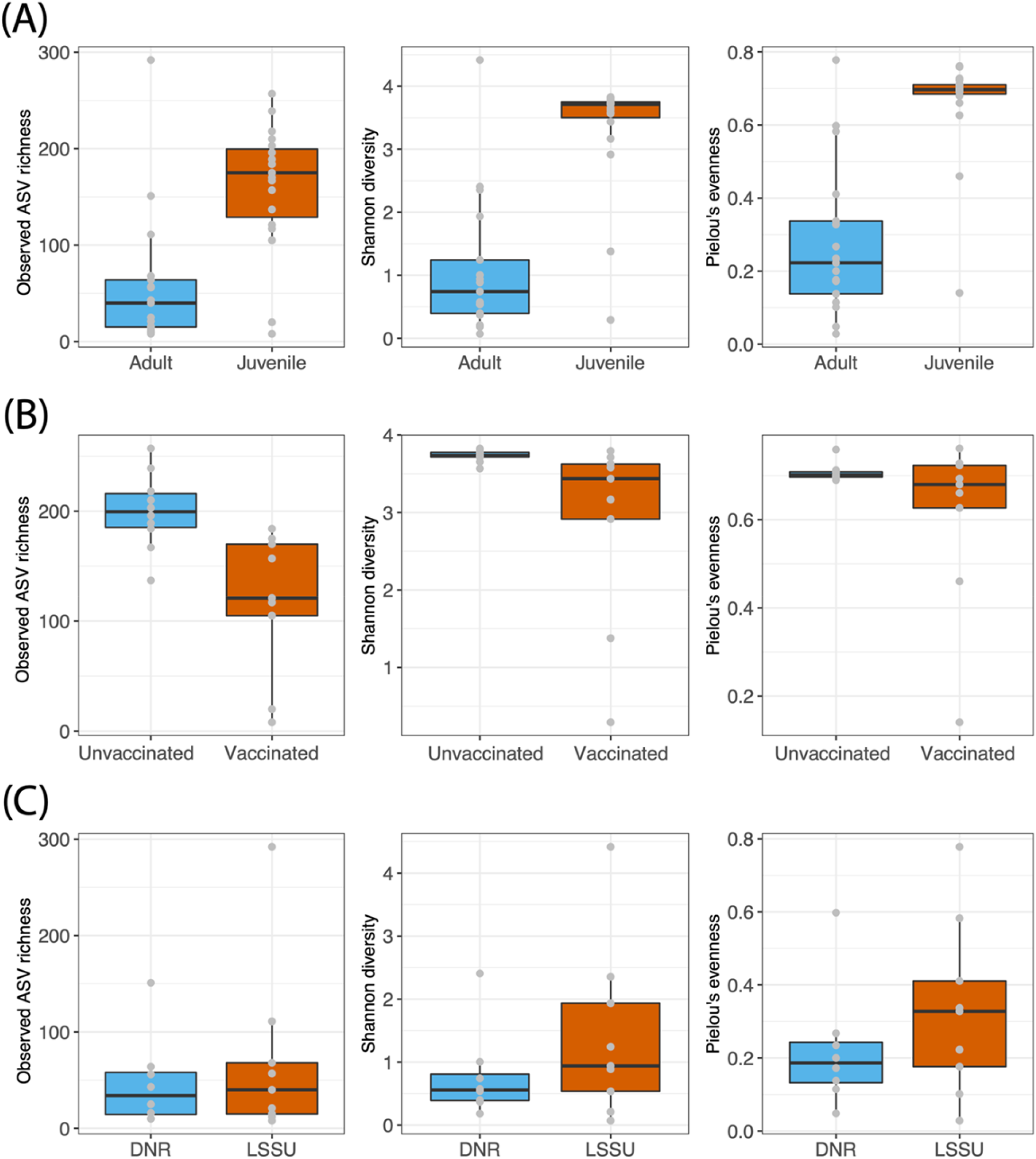
Alpha diversity metrics (observed ASV richness, Shannon index, and Pielou’s evenness) for Atlantic salmon individuals differing in (A) life stage; (B) sub-adult vaccination status; and (C) adult hatchery origin.

### Microbial beta diversity

Principal coordinate analysis of Bray-Curtis distances indicated differences in the structure of sub-adult and adult microbiomes, which displayed differences in their group centroids (PERMANOVA p_adjusted_ < 0.001; Figure 3; Table 2). The dispersion of adult individuals was also higher than sub-adults (PERMDISP p_adjusted_ = 0.001), with an average distance to the centroid of 0.591 for adults and 0.227 for sub-adults. Comparisons of vaccinated and unvaccinated sub-adult individuals also indicated differences in group centroids and dispersion (PERMANOVA p_adjusted_ = 0.001; PERMDISP p_adjusted_ < 0.001; Figure 3; Table 2), with an average distance to the centroid of 0.074 for unvaccinated sub-adults and 0.363 for vaccinated sub-adults. Beta diversity in microbiomes of adult Atlantic salmon did not differ among individuals originating from the LSSU and DNR hatcheries (PERMANOVA p_adjusted_= 0. 865; PERMDISP p_adjusted_ = 0.802; Figure 3; Table 2).

**Table 2.**
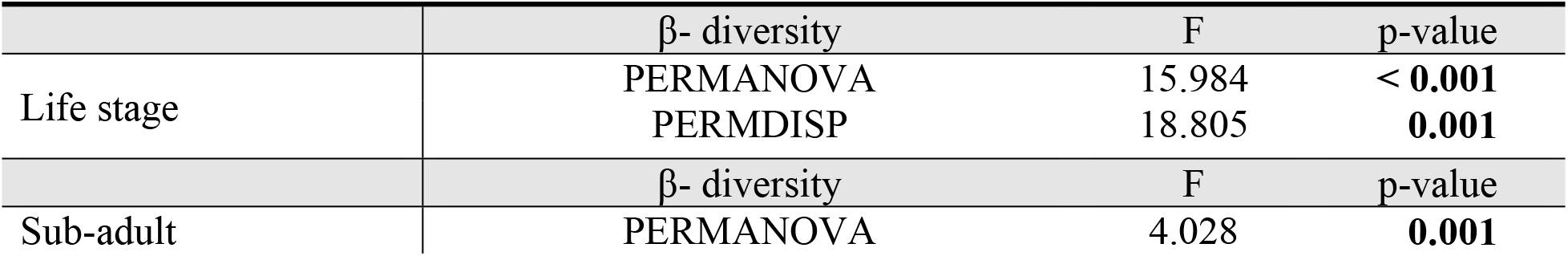
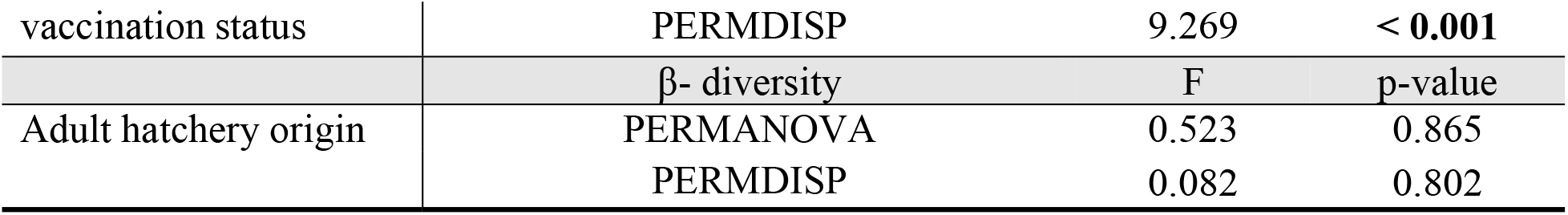
Beta diversity statistics for gut microbiomes of individuals differing in life stage [sub-adults (*n* = 19) vs. adults (*n* = 17)], sub-adult BKD vaccination [unvaccinated (*n* = 10) vs. vaccinated (*n* = 9)], and adult hatchery origin [LSSU (*n* = 9) vs. DNR (*n* = 8)]. Differences in beta diversity (centroid means and dispersion) were evaluated using PERMANOVA and PERMDISP, with p-values calculated from a distribution of 1,000 random permutations and bold values indicating statistical differences at alpha = 0.05.

**Figure 3.**
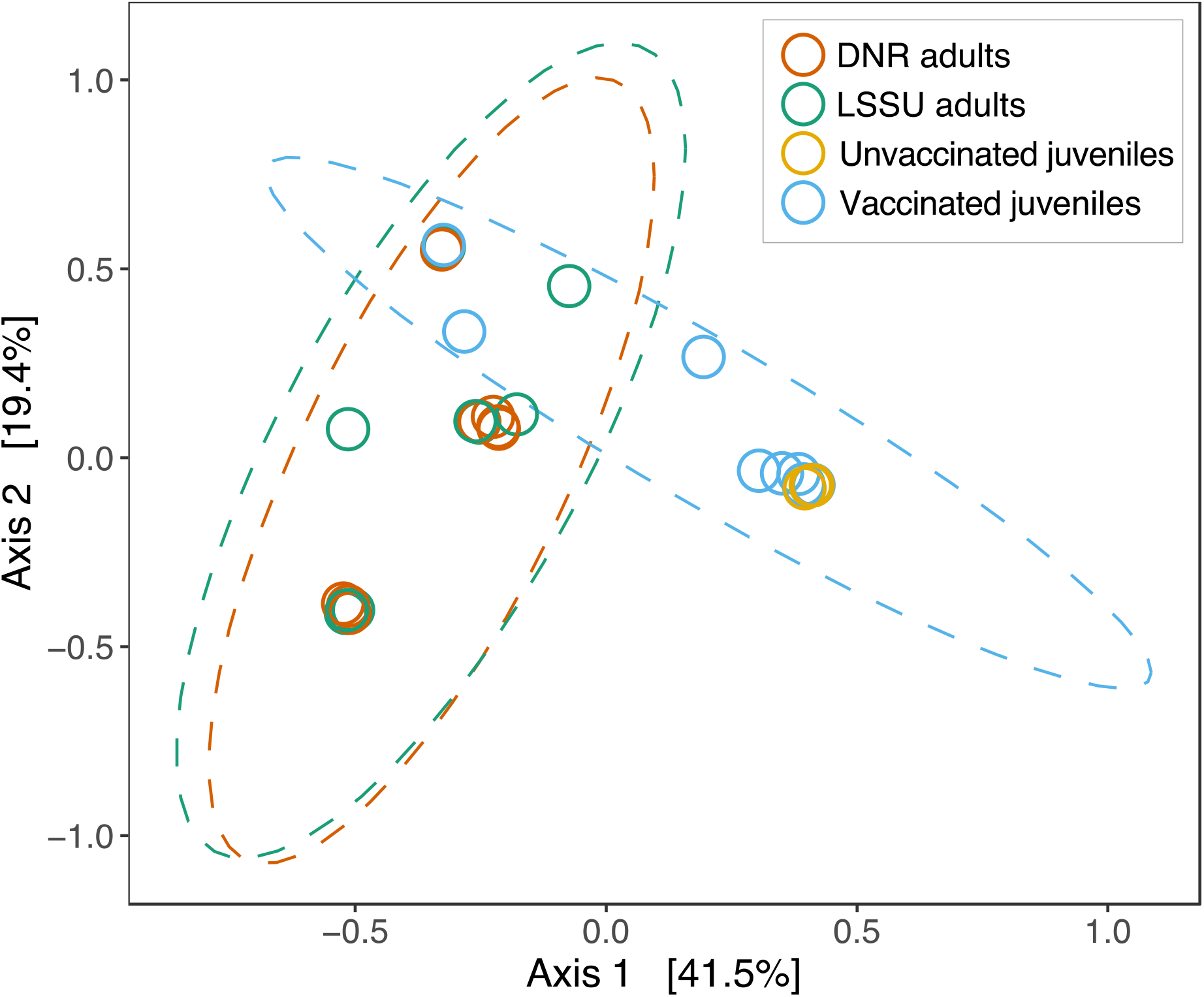
PCoA of Bray-Curtis distances among individual microbiomes shaded based on adult hatchery origin and sub-adult BKD vaccination status. Ellipses represent 95% confidence interval (CI) around the group centroid. Due to low beta diversity among unvaccinated sub-adults, the ellipse representing the 95% CI is not visible.

### Core communities and biomarkers

Adult Atlantic salmon microbiomes did not exhibit highly prevalent microbiota (i.e., taxa present in most of the individuals) at the ASV, genus, or family levels, indicating that the adult salmon gut microbiota did not exhibit a “core-satellite” model (Figure S2, left column). Only *Aeromonadaceae* bacteria were prevalent in 100% of adult fish and an unclassified bacterial family was 88.2% prevalent (Figure 4A). Sub-adult Atlantic salmon, on the other hand, exhibited several highly prevalent microbiota at the ASV, genus, and family levels for both vaccinated and unvaccinated groups (Figure S2, middle and right columns). BKD-vaccinated sub-adults shared 7 core families with ≥ 80% prevalence (Figure 4B) while unvaccinated sub-adults shared 33 core families (Figure 4C). Across all individuals, families that were highly prevalent also tended to have high mean abundance values, but this was not always the case.

**Figure 4.**
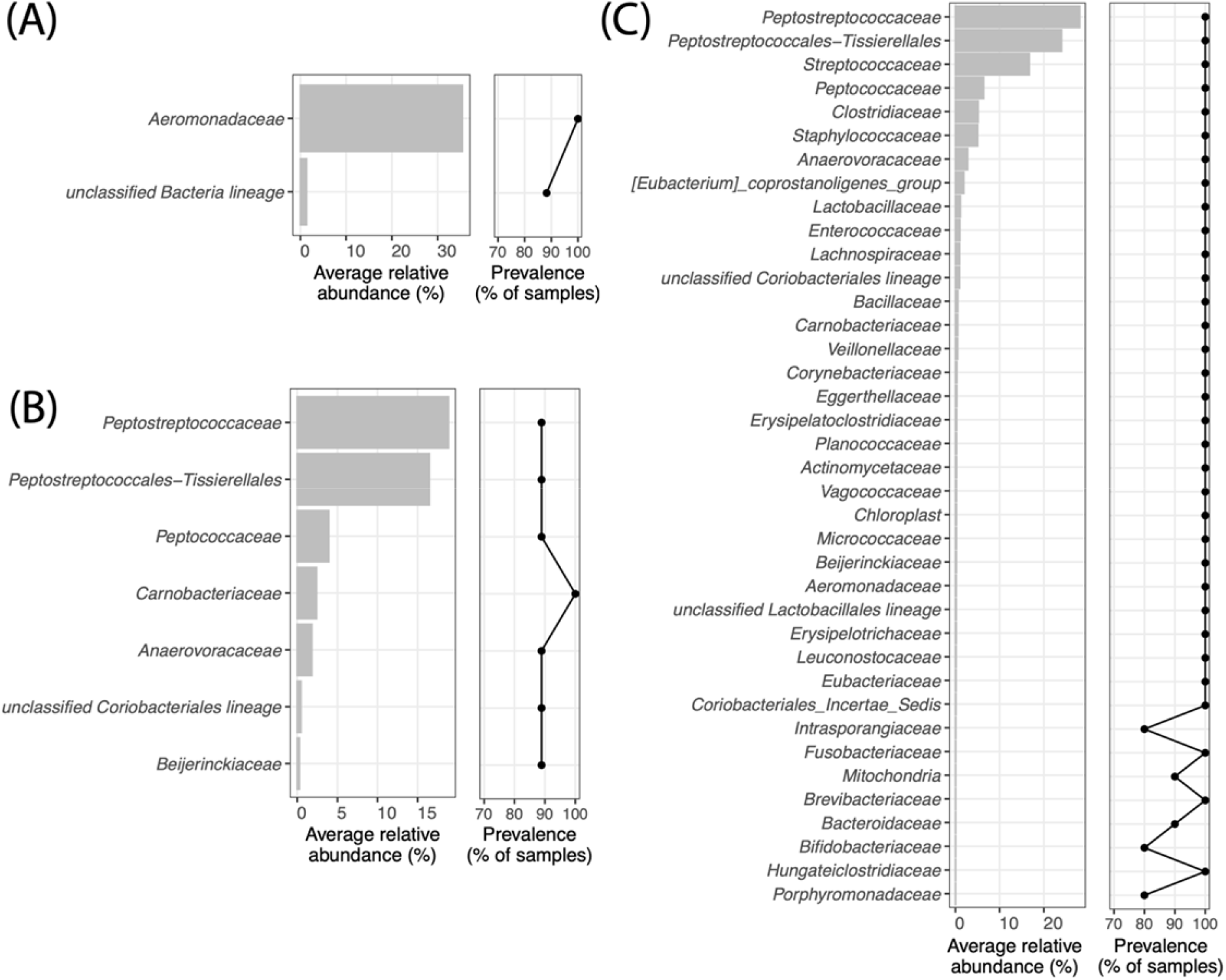
Average relative abundance and prevalence (percentage of samples in which a family is present) for core families of Atlantic salmon microbiomes with ≥ 80% prevalence across (A) all Adults; (B) BKD-vaccinated sub-adults; and (C) Unvaccinated sub-adults.

In the biomarker analysis, we found 12 family-level and 12 genus-level biomarkers across sub-adult and adult Atlantic salmon, with about half of the biomarkers at each level representing each group (Figure 5A). Fewer biomarkers were identified for BKD-vaccinated and unvaccinated sub-adults, with 6 at the family level and 5 and the genus level (Figure 5B).

**Figure 5.**
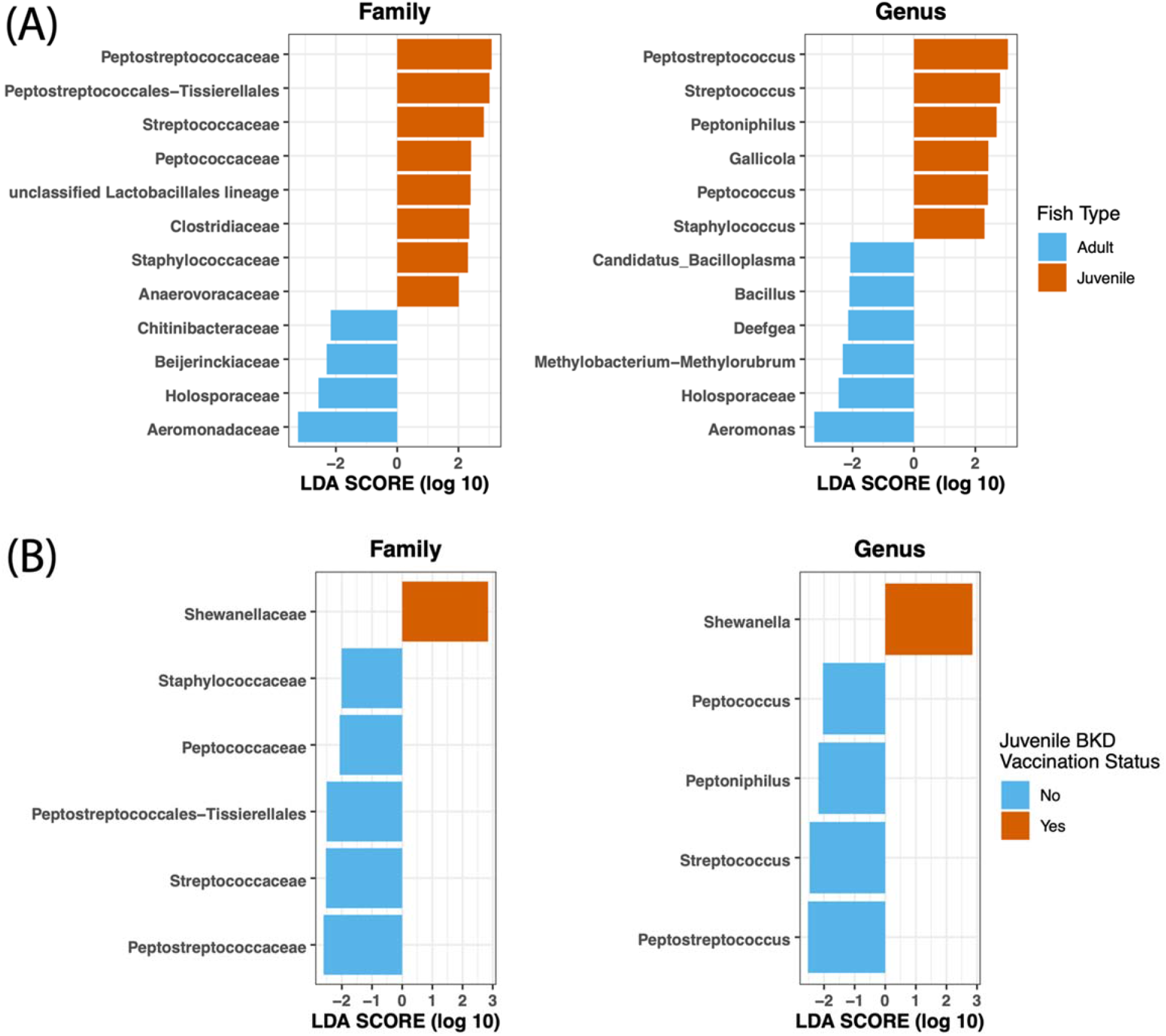
LDA strength of biomarkers in Atlantic salmon gut microbiota at the family and genus levels for individuals grouped by (A) life stage; and (B) sub-adult BKD vaccination status.

## DISCUSSION

The gut microbiome of hatchery-reared Atlantic salmon is influenced by several complex factors, but the exact nature of such factors and effects on host health and physiology remain poorly understood. Thus far, studies investigating fish intestinal microbiomes are relatively limited, especially in relation to understanding the impacts of vaccines on diversity and composition of the gut microbiome. In this study, we found that both the vaccination status and life stage were important factors shaping Atlantic salmon gut microbiota, but hatchery origin was not. Sub-adults had greater gut microbial diversity than adults, while adults displayed greater diversity in gut microbial communities between individuals. BKD vaccination reduced gut microbial diversity within individual salmon and resulted in dissimilar gut microbial communities between individuals. There were no differences in gut microbial communities between adults raised in different hatcheries after they were in the wild for at least one year.

Across all sampled individuals, we found low phylogenetic diversity, with only three phyla (*Firmicutes, Proteobacteria*, and *Actinobacteriota*) comprising over 97% of the sequence reads. This finding is in accordance with previous research documenting high abundances of these phyla in freshwater fishes (1, 33, 53–56). Sub-adult gut microbiomes were dominated by *Firmicutes*, while adult gut microbiomes were dominated by *Proteobacteria*. Previous research investigating differences in the gut microbiomes related to growth rate in Rainbow trout (*Oncorhynchus mykiss*) and common carp (*Cyprinus carpio*) found that fast-growing individuals had a higher proportion of *Firmicutes*, a phylum that may increase fatty acid uptake and impact carbohydrate metabolism in fishes (57–59). Thus, a higher proportion of *Firmicutes* may be associated with fast growth of hatchery-reared sub-adults.

We found that gut microbiomes in sub-adults exhibited significantly higher alpha diversity but reduced beta diversity when compared to adults. While the gut microbiome has been shown to vary throughout the fish life cycle (33, 60–62), sub-adults in this study were collected from the hatchery while adults were captured from the wild. Therefore, recognized differences in gut microbiomes between these two groups may have been impacted by a breadth of covarying factors including host environment, age, and diet. Previous research has shown that wild populations tend to have higher microbial diversity and more specialized gut microbiota than hatchery-reared salmon, which exhibit lower microbial diversity and more generalist bacteria species (21, 32, 63). Uren Webster (63) suggested that because the microbial communities of water in hatcheries are typically impoverished compared to natural environments, reduced microbiome diversity in hatcheries may be the result of a limited microbial species pool in the immediate environment. However, because the water for the LSSU hatchery is sourced from unfiltered, untreated water from the St. Marys River, sub-adults and adults in this study would have been experiencing similar microbial communities in the surrounding environment. Gut microbiomes may therefore not be strongly impacted by the microbes in the surrounding environment but instead by other environmental factors, host age, and/or diet.

In a previous study, the lowest levels of alpha diversity in wild Atlantic salmon were observed in later stages of life (33), possibly attributed to dietary complexity in juvenile salmonids (64). In this study, however, sub-adults at the hatchery were fed commercial fish pellet food (Bio-Oregon and Skretting), and adults in the wild likely would have been consuming a variety of fish and invertebrates as they returned to the river to spawn. In addition to the complexity and availability of dietary items, hatchery and wild environments differ in many other ways including the density of individuals, environmental heterogeneity, and intra- and interspecific interactions, all of which could impact the diversity and composition of gut microbiomes. A breadth of research supports the importance of both host diet and environment in determining the gut microbiome of fishes (1, 55, 63, 65–69), and because these factors covary in our experimental design, we attribute differences in microbiome diversity among life stages to be driven by differences in both habitat and diet.

In addition to reduced alpha diversity, we found that adult Atlantic salmon exhibited greater variation among individuals (beta diversity) in their microbial communities compared to sub-adults. This is in contrast to previous research showing that neutral processes generate a substantial amount of variation in sub-adult fish gut microbial communities, but that the relative importance of non-neutral processes (e.g., microbe-microbe interactions, dispersal, or host selection) increases with host age, resulting in more similar gut microbiomes at later life stages (70). As highlighted above, the differences we observed among sub-adults and adults may therefore be more strongly impacted by host environment and diet than age. Wild populations exhibit more specialized gut microbiota, whereas captive-reared salmon hosted more generalist bacteria, thereby resulting in increased similarity in communities across individuals (21). In addition to increased environmental heterogeneity experienced by wild adults, Atlantic salmon are opportunistic generalists and will readily consume a variety of foods (71, 72). Therefore, enhanced beta diversity in the gut microbiome of adult salmon in our study may be due to increased variability in environmental and dietary conditions experienced in the wild compared to the hatchery. Furthermore, hatcheries are a much more densely populated environment compared to natural systems, increasing the likelihood of similar colonization of microbes between hatchery fish (21).

Previous research has demonstrated that the administration of antibiotics can disrupt microbial communities (73), most of which focuses on how the microbiome may affect vaccine efficacy (27). However, it is not well understood how vaccines influence gut microbial communities. We found that BKD vaccination in sub-adult salmon greatly reduced gut microbial ASV richness and alpha diversity. In turn, this reduction in diversity resulted in a large increase in beta diversity and dispersion while also erasing any trace of a robust core microbiome. In unvaccinated sub-adults, we identified 33 core families, but in vaccinated sub-adults we only identified seven core families. Because the identity and diversity of early microbial colonizers can affect establishment of future assemblages (31, 63), the effects of BKD vaccination could have important downstream consequences for Atlantic salmon health and physiology. For instance, variation in gut microbiota may impact hosts’ ability to prevent infection by pathogenic microbes (74) or adapt to novel environmental conditions through phenotypic plasticity (75).

Given the importance of protecting salmon from BKD infection, determining the consequences of this vaccine on microbiome composition and function is an important next step in designing solutions to support host health. For instance, supplementing hosts with pre- or probiotics could mitigate the loss of important bacterial taxa during the vaccine inoculation process.

Previous work has shown that hatchery rearing conditions such as water circulation, water treatment, and diet shape microbial composition of the individuals being raised (21, 24, 25, 32, 63). However, we did not detect any significant differences in the structure or diversity of gut microbiota of adult fish that were initially reared in the LSSU and DNR hatcheries, despite differences in water sources at these facilities (i.e., ambient river water at LSSU vs. filtered, treated pond water at DNR). This could be due to the fact that all gametes collected for artificial spawning were initially sourced from the same broodstock and fertilized at LSSU before a portion of the fertilized eggs were transferred to the DNR hatchery. Microbial colonization in the fish GI tract begins at the earliest life stages, when the effects of the regional microbial species pool in the surrounding environment exhibit the strongest effects on fish microbiomes (76, 77). Thus, by the time the fertilized eggs were transferred to the DNR hatchery, community assembly of fish gut microbiomes may have been less impacted by the particular hatchery conditions.

Atlantic salmon were also fed the same commercial pellet food at both hatcheries, so differences among hatcheries may not be observed if the influence of diet on the gut microbiome outweighs the importance of environment.

An alternative explanation for the lack of differences in the gut microbiome among Atlantic salmon adults sourced from different hatcheries could be that the influence of hatchery origin disappeared after individuals were released into the wild. Adult individuals had experienced the open water of the St. Marys River, and most likely Lake Huron, for at least one year before being recaptured and assessed; any fish that had emigrated to Lake Huron would have returned to the St. Marys River and staged for several weeks or months prior to capture.

Although the early gut microbiome has been shown to impact gut microbial assemblages later in life (78), the microbiome in Atlantic salmon is also extremely plastic (32) and legacy effects have been shown to gradually disappear after experiencing identical environmental conditions (61). This could also indicate that the differences we observed in microbiomes of BKD-vaccinated and unvaccinated sub-adults will converge once the fish are released into the wild.

More research is needed to better understand the legacy effects of vaccination status on the microbiome of adults that have been living in a common environment.

Acquiring a basic understanding of the ecological processes that govern microbial assemblages like the gut microbiome in Atlantic salmon is important for developing strategies to improve the rearing and reintroduction success of captive-reared fishes. By identifying the specific microbiota, understanding how gut microbial communities assemble, and elucidating the functional role of gut microbes in the health of salmon, fisheries managers can start to implement practices, such as prebiotic or antibiotic treatments, that may reduce operational costs and increase survival. Further research linking the composition of the gut microbiome to specific functions and investigating the impacts of such function on host health will be essential for predicting and possibly manipulating the microbiome to improve performance and health in fishes (28, 79). However, this research provides foundational knowledge on the diversity and structure of gut microbiomes among different groups of Atlantic salmon, providing a basis for inferences about the importance of life stage and vaccination status on gut microbiomes in hatchery-reared fishes.

## ACKNOWLEDGEMENTS

We thank Roger Greil (LSSU Center for Freshwater Research and Education Fish Hatchery), Ed Eisch (Michigan Department of Natural Resources), and many hatchery personnel who contributed to hatchery operations that provided fish for our study. Funding for this research was supported by Rapid Funds from the Cooperative Institute of Great Lakes Research awarded to KK, AM, JK, and FL and a Living Earth Collaborative Seed Grant awarded to JK and FL. KJA is supported by a Postdoctoral Research Fellowship from the Living Earth Collaborative. LEJ is supported by the National Science Foundation grant DGE-2139839.

## DATA ACCESSIBILITY

All raw sequence data from this experiment were submitted to the NCBI Sequence Read Archive under BioProject ID PRJNA932862. Source code including bioinformatic processing and statistical analysis is available through https://github.com/linglab-washu/salmon_microbiome.

## REFERENCES

1. Llewellyn MS, Boutin S, Hoseinifar SH, Derome N. 2014. Teleost microbiomes: the state of the art in their characterization, manipulation and importance in aquaculture and fisheries. Frontiers in Microbiology 5:207.

2. McFall-Ngai M, Hadfield MG, Bosch TC, Carey HV, Domazet-Lošo T, Douglas AE, Dubilier N, Eberl G, Fukami T, Gilbert SF. 2013. Animals in a bacterial world, a new imperative for the life sciences. Proceedings of the National Academy of Sciences 110:3229–3236.

3. Nayak SK. 2010. Role of gastrointestinal microbiota in fish. Aquaculture Research 41:1553–1573.

4. Suzuki TA. 2017. Links between natural variation in the microbiome and host fitness in wild mammals. Integrative and Comparative Biology 57:756–769.

5. West AG, Waite DW, Deines P, Bourne DG, Digby A, McKenzie VJ, Taylor MW. 2019. The microbiome in threatened species conservation. Biological Conservation 229:85–98.

6. Bahrndorff S, Alemu T, Alemneh T, Lund Nielsen J. 2016. The Microbiome of Animals: Implications for Conservation Biology. International Journal of Genomics 2016:e5304028.

7. Jiménez RR, Sommer S. 2017. The amphibian microbiome: natural range of variation, pathogenic dysbiosis, and role in conservation. Biodiversity and Conservation 26:763–786.

8. Stumpf RM, Gomez A, Amato KR, Yeoman CJ, Polk JD, Wilson BA, Nelson KE, White BA, Leigh SR. 2016. Microbiomes, metagenomics, and primate conservation: New strategies, tools, and applications. Biological Conservation 199:56–66.

9. Yukgehnaish K, Kumar P, Sivachandran P, Marimuthu K, Arshad A, Paray BA, Arockiaraj J. 2020. Gut microbiota metagenomics in aquaculture: factors influencing gut microbiome and its physiological role in fish. Reviews in Aquaculture 12:1903–1927.

10. Langlois L, Akhtar N, Tam KC, Dixon B, Reid G. 2021. Fishing for the right probiotic: host–microbe interactions at the interface of effective aquaculture strategies. FEMS Microbiology Reviews 45:fuab030.

11. Ramos M, Gonçalves J, Batista S, Costas B, Pires M, Rema P, Ozório R. 2015. Growth, immune responses and intestinal morphology of rainbow trout (Oncorhynchus mykiss) supplemented with commercial probiotics. Fish & Shellfish Immunology 45:19–26.

12. Wang J, Ji H, Wang S, Liu H, Zhang W, Zhang D, Wang Y. 2018. Probiotic Lactobacillus plantarum promotes intestinal barrier function by strengthening the epithelium and modulating gut microbiota. Frontiers in Microbiology 9:1953.

13. Bunnell DB, Barbiero RP, Ludsin SA, Madenjian CP, Warren GJ, Dolan DM, Brenden TO, Briland R, Gorman OT, He JX. 2014. Changing ecosystem dynamics in the Laurentian Great Lakes: bottom-up and top-down regulation. Bioscience 64:26–39.

14. Dymond JR, MacKay HH, Burridge ME, Holm E, Bird PW. 2019. The history of the Atlantic salmon in Lake Ontario. Aquatic Ecosystem Health & Management 22:305–315.

15. Parsons JW. 1973. History of salmon in the Great Lakes, 1850-1970. Federal Government Series. U.S. Bureau of Sport Fisheries and Wildlife.

16. Tucker S, Moerke A, Steinhart G, Greil R. 2014. NOTE: First record of natural reproduction by Atlantic salmon (Salmo salar) in the St. Marys River, Michigan. Journal of Great Lakes Research 40:1022–1026.

17. Behmer DJ, Greil RW, Scott SJ, Hanna T. 1993. Harvest and movement of Atlantic salmon stocked in the St. Marys River, Michigan. Journal of Great Lakes Research 19:533–540.

18. Crawford SS, Canada NRC. 2001. Salmonine Introductions to the Laurentian Great Lakes: An Historical Review and Evaluation of Ecological Effects. NRC Research Press.

19. Scott RJ, Noakes DLG, Beamish FWH, Carl LM. 2003. Chinook salmon impede Atlantic salmon conservation in Lake Ontario. Ecology of Freshwater Fish 12:66–73.

20. Stringwell R, Lock A, Stutchbury C, Baggett E, Taylor J, Gough P, Garcia de Leaniz C. 2014. Maladaptation and phenotypic mismatch in hatchery-reared Atlantic salmon Salmo salar released in the wild. Journal of Fish Biology 85:1927–1945.

21. Lavoie C, Courcelle M, Redivo B, Derome N. 2018. Structural and compositional mismatch between captive and wild Atlantic salmon (Salmo salar) parrs’ gut microbiota highlights the relevance of integrating molecular ecology for management and conservation methods. Evolutionary Applications 11:1671–1685.

22. Lavoie C, Wellband K, Perreault A, Bernatchez L, Derome N. 2021. Artificial Rearing of Atlantic Salmon Juveniles for Supportive Breeding Programs Induces Long-Term Effects on Gut Microbiota after Stocking. 9. Microorganisms 9:1932.

23. Walter J, Ley R. 2011. The Human Gut Microbiome: Ecology and Recent Evolutionary Changes. Annual Review of Microbiology 65:411–429.

24. Dehler CE, Secombes CJ, Martin SAM. 2017. Environmental and physiological factors shape the gut microbiota of Atlantic salmon parr (Salmo salar L.). Aquaculture 467:149–157.

25. Minich JJ, Poore GD, Jantawongsri K, Johnston C, Bowie K, Bowman J, Knight R, Nowak B, Allen EE. 2020. Microbial Ecology of Atlantic Salmon (Salmo salar) Hatcheries: Impacts of the Built Environment on Fish Mucosal Microbiota. Applied and Environmental Microbiology 86:e00411–20.

26. Liu L, Gong Y-X, Zhu B, Liu G-L, Wang G-X, Ling F. 2015. Effect of a new recombinant Aeromonas hydrophila vaccine on the grass carp intestinal microbiota and correlations with immunological responses. Fish & Shellfish Immunology 45:175–183.

27. Di Renzo L, Franza L, Monsignore D, Esposito E, Rio P, Gasbarrini A, Gambassi G, Cianci R, De Lorenzo A. 2022. Vaccines, Microbiota and Immunonutrition: Food for Thought. Vaccines 10:294.

28. Perry WB, Lindsay E, Payne CJ, Brodie C, Kazlauskaite R. 2020. The role of the gut microbiome in sustainable teleost aquaculture. Proceedings of the Royal Society B: Biological Sciences 287:20200184.

29. Ringø E, Zhou Z, Vecino J l. g., Wadsworth S, Romero J, Krogdahl Å, Olsen R e., Dimitroglou A, Foey A, Davies S, Owen M, Lauzon H l., Martinsen L l., De Schryver P, Bossier P, Sperstad S, Merrifield D l. 2016. Effect of dietary components on the gut microbiota of aquatic animals. A never-ending story? Aquaculture Nutrition 22:219–282.

30. Gensollen T, Iyer SS, Kasper DL, Blumberg RS. 2016. How colonization by microbiota in early life shapes the immune system. Science 352:539–544.

31. Martínez I, Maldonado-Gomez MX, Gomes-Neto JC, Kittana H, Ding H, Schmaltz R, Joglekar P, Cardona RJ, Marsteller NL, Kembel SW, Benson AK, Peterson DA, Ramer-Tait AE, Walter J. 2018. Experimental evaluation of the importance of colonization history in early-life gut microbiota assembly. eLife 7:e36521.

32. Uren Webster TM, Rodriguez-Barreto D, Castaldo G, Gough P, Consuegra S, Garcia de Leaniz C. 2020. Environmental plasticity and colonisation history in the Atlantic salmon microbiome: A translocation experiment. Molecular Ecology 29:886–898.

33. Llewellyn MS, McGinnity P, Dionne M, Letourneau J, Thonier F, Carvalho GR, Creer S, Derome N. 2016. The biogeography of the atlantic salmon (Salmo salar) gut microbiome. 5. The ISME Journal 10:1280–1284.

34. Elliott DG, Wiens GD, Hammell KL, Rhodes LD. 2014. Vaccination against Bacterial Kidney Disease, p. 255–272. In Fish Vaccination. John Wiley & Sons, Ltd.

35. Evelyn TPT. 1993. Bacterial kidney disease – BKD, p. 177–195. In Inglis, V, Roberts, RJ, Bromage, NR (eds.), Bacterial Diseases of Fish. Halsted Press, New York.

36. Salonius K, Siderakis C, MacKinnon AM, Griffiths SG. 2005. Use of Arthrobacter davidanieli as a live vaccine against Renibacterium salmoninarum and Piscirickettsia salmonis in salmonids. Developments in Biologicals 121:189–197.

37. Schmidt TM, DeLong EF, Pace NR. 1991. Analysis of a marine picoplankton community by 16S rRNA gene cloning and sequencing. Journal of Bacteriology 173:4371–4378.

38. Preheim SP, Perrotta AR, Friedman J, Smilie C, Brito I, Smith MB, Alm E. 2013. Chapter Eighteen - Computational Methods for High-Throughput Comparative Analyses of Natural Microbial Communities, p. 353–370. In DeLong, EF (ed.), Methods in Enzymology. Academic Press.

39. Caporaso JG, Lauber CL, Walters WA, Berg-Lyons D, Lozupone CA, Turnbaugh PJ, Fierer N, Knight R. 2011. Global patterns of 16S rRNA diversity at a depth of millions of sequences per sample. Proceedings of the National Academy of Sciences 108:4516–4522.

40. Bolyen E, Rideout JR, Dillon MR, Bokulich NA, Abnet CC, Al-Ghalith GA, Alexander H, Alm EJ, Arumugam M, Asnicar F. 2019. Reproducible, interactive, scalable and extensible microbiome data science using QIIME 2. Nature Biotechnology 37:852–857.

41. Callahan BJ, McMurdie PJ, Rosen MJ, Han AW, Johnson AJA, Holmes SP. 2016. DADA2: high-resolution sample inference from Illumina amplicon data. Nature Methods 13:581–583.

42. Bokulich NA, Kaehler BD, Rideout JR, Dillon M, Bolyen E, Knight R, Huttley GA, Gregory Caporaso J. 2018. Optimizing taxonomic classification of marker-gene amplicon sequences with QIIME 2’s q2-feature-classifier plugin. Microbiome 6:1–17.

43. Quast C, Pruesse E, Yilmaz P, Gerken J, Schweer T, Yarza P, Peplies J, Glöckner FO. 2012. The SILVA ribosomal RNA gene database project: improved data processing and web-based tools. Nucleic Acids Research 41:D590–D596.

44. Bisanz J. 2022. Tutorial: Integrating QIIME2 and R for data visualization and analysis using qiime2R (v0.99.6). R.

45. McMurdie PJ, Holmes S. 2013. phyloseq: an R package for reproducible interactive analysis and graphics of microbiome census data. PloS one 8:e61217.

46. Davis NM, Proctor DM, Holmes SP, Relman DA, Callahan BJ. 2018. Simple statistical identification and removal of contaminant sequences in marker-gene and metagenomics data. Microbiome 6:1–14.

47. R Core Team. 2021. R: A language and environment for statistical computing. R Foundation for Statistical Computing, Vienna, Austria.

48. Wickham H, Chang W, Wickham MH. 2016. Package ‘ggplot2.’ Create elegant data visualisations using the grammar of graphics Version 2:1–189.

49. Oksanen J, Blanchet FG, Kindt R, Legendre P, Minchin PR, O’hara R, Simpson GL, Solymos P, Stevens MHH, Wagner H. 2013. Package ‘vegan.’ Community Ecology Package, version 2:1–295.

50. Paradis E, Claude J, Strimmer K. 2004. APE: analyses of phylogenetics and evolution in R language. Bioinformatics 20:289–290.

51. Kassambara A. 2020. rstatix: Pipe-friendly framework for basic statistical tests. R package version 06 0.

52. Segata N, Izard J, Waldron L, Gevers D, Miropolsky L, Garrett WS, Huttenhower C. 2011. Metagenomic biomarker discovery and explanation. Genome Biology 12:1–18.

53. Gómez GD, Balcázar JL. 2008. A review on the interactions between gut microbiota and innate immunity of fish. FEMS Immunology & Medical Microbiology 52:145–154.

54. Roeselers G, Mittge EK, Stephens WZ, Parichy DM, Cavanaugh CM, Guillemin K, Rawls JF. 2011. Evidence for a core gut microbiota in the zebrafish. The ISME Journal 5:1595–1608.

55. Sullam KE, Essinger SD, Lozupone CA, O’CONNOR MP, Rosen GL, Knight R, Kilham SS, Russell JA. 2012. Environmental and ecological factors that shape the gut bacterial communities of fish: a meta-analysis. Molecular Ecology 21:3363–3378.

56. Wu S, Wang G, Angert ER, Wang W, Li W, Zou H. 2012. Composition, diversity, and origin of the bacterial community in grass carp intestine. PloS One 7:e30440.

57. Chapagain P, Arivett B, Cleveland BM, Walker DM, Salem M. 2019. Analysis of the fecal microbiota of fast-and slow-growing rainbow trout (Oncorhynchus mykiss). BMC genomics 20:1–11.

58. Li X, Yan Q, Xie S, Hu W, Yu Y, Hu Z. 2013. Gut Microbiota Contributes to the Growth of Fast-Growing Transgenic Common Carp (Cyprinus carpio L.). PLoS One 8:e64577.

59. Semova I, Carten JD, Stombaugh J, Mackey LC, Knight R, Farber SA, Rawls JF. 2012. Microbiota regulate intestinal absorption and metabolism of fatty acids in the zebrafish. Cell Host & Microbe 12:277–288.

60. Bledsoe JW, Peterson BC, Swanson KS, Small BC. 2016. Ontogenetic characterization of the intestinal microbiota of channel catfish through 16S rRNA gene sequencing reveals insights on temporal shifts and the influence of environmental microbes. PloS one 11:e0166379.

61. Deng Y, Kokou F, Eding EH, Verdegem MCJ. 2021. Impact of early-life rearing history on gut microbiome succession and performance of Nile tilapia. Animal Microbiome 3:81.

62. Zhang Z, Li D, Refaey MM, Xu W, Tang R, Li L. 2018. Host age affects the development of southern catfish gut bacterial community divergent from that in the food and rearing water. Frontiers in Microbiology 9:495.

63. Uren Webster TM, Consuegra S, Hitchings M, Garcia de Leaniz C. 2018. Interpopulation Variation in the Atlantic Salmon Microbiome Reflects Environmental and Genetic Diversity. Applied and Environmental Microbiology 84:e00691–18.

64. Orlov AV, Gerasimov YV, Lapshin OM. 2006. The feeding behaviour of cultured and wild Atlantic salmon, Salmo salar L., in the Louvenga River, Kola Peninsula, Russia. ICES Journal of Marine Science 63:1297–1303.

65. Kim PS, Shin N-R, Lee J-B, Kim M-S, Whon TW, Hyun D-W, Yun J-H, Jung M-J, Kim JY, Bae J-W. 2021. Host habitat is the major determinant of the gut microbiome of fish. Microbiome 9:166.

66. Lokesh J, Kiron V. 2016. Transition from freshwater to seawater reshapes the skin-associated microbiota of Atlantic salmon. 1. Scientific Reports 6:19707.

67. Kashinskaya E n., Simonov E p., Kabilov M r., Izvekova G i., Andree K b., Solovyev M m. 2018. Diet and other environmental factors shape the bacterial communities of fish gut in an eutrophic lake. Journal of Applied Microbiology 125:1626–1641.

68. Ingerslev H-C, von Gersdorff Jørgensen L, Lenz Strube M, Larsen N, Dalsgaard I, Boye M, Madsen L. 2014. The development of the gut microbiota in rainbow trout (Oncorhynchus mykiss) is affected by first feeding and diet type. Aquaculture 424–425:24–34.

69. Smith CC, Snowberg LK, Gregory Caporaso J, Knight R, Bolnick DI. 2015. Dietary input of microbes and host genetic variation shape among-population differences in stickleback gut microbiota. 11. The ISME Journal 9:2515–2526.

70. Burns AR, Stephens WZ, Stagaman K, Wong S, Rawls JF, Guillemin K, Bohannan BJ. 2016. Contribution of neutral processes to the assembly of gut microbial communities in the zebrafish over host development. The ISME journal 10:655–664.

71. Dixon HJ, Dempson JB, Sheehan TF, Renkawitz MD, Power M. 2017. Assessing the diet of North American Atlantic salmon (Salmo salar L.) off the West Greenland coast using gut content and stable isotope analyses. Fisheries Oceanography 26:555–568.

72. Roseman EF, Schaeffer JS, Bright E, Fielder DG. 2014. Angler-Caught Piscivore Diets Reflect Fish Community Changes in Lake Huron. Transactions of the American Fisheries Society 143:1419–1433.

73. Parra M, Espinoza D, Valdes N, Vargas R, Gonzalez A, Modak B, Tello M. 2020. Microbiota Modulates the Immunomodulatory Effects of Filifolinone on Atlantic Salmon. Microorganisms 8:1320.

74. Li T, Long M, Ji C, Shen Z, Gatesoupe F-J, Zhang X, Zhang Q, Zhang L, Zhao Y, Liu X. 2016. Alterations of the gut microbiome of largemouth bronze gudgeon (Coreius guichenoti) suffering from furunculosis. Scientific Reports 6:1–9.

75. Alberdi A, Aizpurua O, Bohmann K, Zepeda-Mendoza ML, Gilbert MTP. 2016. Do Vertebrate Gut Metagenomes Confer Rapid Ecological Adaptation? Trends in Ecology & Evolution 31:689–699.

76. Stephens WZ, Burns AR, Stagaman K, Wong S, Rawls JF, Guillemin K, Bohannan BJ. 2016. The composition of the zebrafish intestinal microbial community varies across development. The ISME Journal 10:644–654.

77. Yan Q, Li J, Yu Y, Wang J, He Z, Van Nostrand JD, Kempher ML, Wu L, Wang Y, Liao L. 2016. Environmental filtering decreases with fish development for the assembly of gut microbiota. Environmental Microbiology 18:4739–4754.

78. Sprockett D, Fukami T, Relman DA. 2018. Role of priority effects in the early-life assembly of the gut microbiota. Nature Reviews Gastroenterology & Hepatology 15:197–205.

79. Talwar C, Nagar S, Lal R, Negi RK. 2018. Fish gut microbiome: current approaches and future perspectives. Indian journal of microbiology 58:397–414.

